# StringTie3 Improves Total RNA-seq Assembly by Resolving Nascent and Mature Transcripts

**DOI:** 10.1101/2025.05.21.655404

**Authors:** Ida Shinder, Geo Pertea, Richard Hu, Zoe Rudnick, Mihaela Pertea

## Abstract

Accurate assembly of rRNA-depleted (total) RNA-seq remains challenging because existing methods often conflate incomplete, nascent RNA with fully processed mature isoforms, leading to misassemblies and quantification errors that skew downstream analyses. Here, we present StringTie3, a major update to the widely used StringTie assembler, specifically designed for total RNA-seq. This new version introduces two key innovations: (1) a nascent mode that models co-transcriptional splicing to separate nascent from mature transcripts, and (2) a refined long-read module that distinguishes genuine polyadenylation sites from poly(A)-priming artifacts. Across short-, long-, and hybrid-read datasets, StringTie3 substantially reduces assembly errors and outperforms existing tools, boosting precision by up to 20% in short-read total RNA-seq and improving sensitivity and precision by as much as 37% and 75%, respectively, in long-read assemblies. In Argonaute knockout experiments, nascent-mode analysis shows that single knockouts predominantly alter nascent transcripts while leaving mature RNA largely unchanged, whereas double or triple knockouts disrupt both fractions. Applying this approach to breast cancer samples shows that, although nascent and mature RNA levels often correlate, certain extracellular matrix and tumor suppressor genes deviate from this pattern, suggesting post-transcriptional regulation. By accurately reconstructing transcriptomes and distinguishing nascent from mature RNA, StringTie3 reveals hidden layers of RNA regulation and provides a powerful framework for investigating transcriptional and post-transcriptional processes in total RNA-seq data.

## Introduction

RNA sequencing (RNA-seq) has substantially advanced our understanding of gene expression and splicing by enabling transcriptional profiling across diverse biological contexts^1^. Early applications primarily relied on poly(A)-selected libraries to enrich for mature messenger RNAs (mRNAs), overlooking non-polyadenylated and nascent transcripts. Over time, RNA-seq has evolved into a versatile toolkit, including single-cell analyses that reveal cellular heterogeneity and specialized methods for nascent transcription^1, 2^. While these approaches have expanded the scope of RNA-seq, most workflows continue to rely on poly(A) selection for transcript assembly and quantification, thus excluding non-polyadenylated RNA–such as histone-encoded transcripts, various non-coding RNAs, and partially processed, nascent RNA. Although specialized nascent RNA assays (e.g., GRO-seq, PRO-seq, 4sU-seq) can capture newly synthesized molecules, they are time- and labor-intensive and require actively transcribing cells, making them impractical for many clinical samples, including biopsies^3^.

Unlike poly(A)-selected methods, total RNA-seq with rRNA depletion preserves both polyadenylated and non-polyadenylated species^4, 5^ (Fig. 1A). Although most transcript assemblers include strategies to handle pre-mRNA contamination, they are less accurate when nascent RNA is abundant. In loci with high nascent expression, partially spliced introns are frequently misassembled as exons (Fig. 1B), obscuring full-length isoforms and confounding downstream analyses such as differential expression. Because nascent transcripts can represent a large share of total RNA, particularly in cells undergoing rapid proliferation or differentiation^4^, ignoring them can obscure essential regulatory events, distort transcriptional dynamics, and introduce quantification biases^6^. In addition, poly(A) selection often enriches for shorter isoforms in degraded or clinical samples^7^, further limiting the effectiveness of conventional methods for accurately reconstructing the transcriptome.

**Figure 1.**
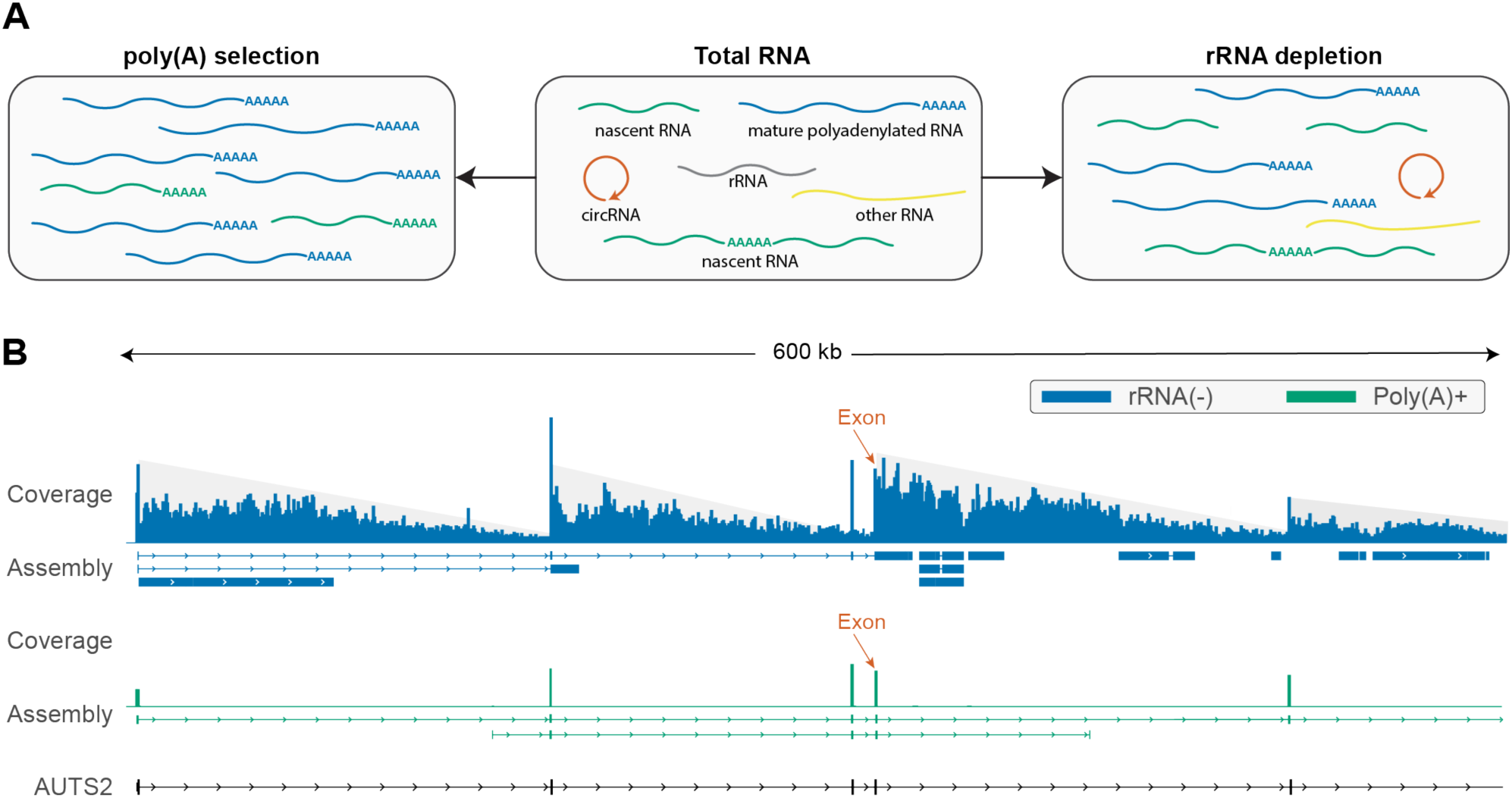
Comparison of poly(A)+ selection and rRNA-depletion reveals misassemblies arising from nascent transcripts. **(A)** Schematic comparing poly(A)+ selection, which enriches for mature polyadenylated RNAs (blue) but may also capture poly(A)-priming artifacts (green transcripts with internal poly(A) sequences), and rRNA depletion (rRNA(-)), which removes ribosomal RNAs (gray) while retaining mature polyadenylated RNAs (blue), non-polyadenylated RNAs (e.g., circRNAs in orange, histone and other RNAs in yellow), and incomplete nascent transcripts (green). **(B)** Read coverage and transcript assemblies at the *AUTS2* locus in rRNA-depleted (blue) and poly(A)+ (green) RNA-seq data, each assembled with StringTie2. The reference MANE annotation^13^ (black) depicts exons (rectangles) and introns (connecting lines). In rRNA-depleted data, intronic reads are misassembled as exons, obscuring an annotated exon (orange arrow) that is accurately reconstructed in poly(A)+ data. The sawtooth pattern (gray highlight) in rRNA-depleted coverage reflects co-transcriptional intron removal.

Here, we present StringTie3, a major update to the widely used StringTie^8, 9^ assembler, designed to explicitly model co-transcriptional splicing in order to distinguish nascent transcripts from mature isoforms in rRNA-depleted libraries. Because most splicing (up to 70%) occurs during transcription rather than afterward^10^, nascent RNA often contains partially spliced introns, resulting in characteristic sawtooth intronic coverage patterns (Fig. 1B and Fig. 2A). These patterns arise because the intron is incrementally transcribed while co-transcriptional splicing progressively removes already transcribed segments, creating overlapping partially spliced intermediates with decreasing read coverage along the intron (Fig. 2A). Building on this insight, StringTie3’s new “nascent mode” prevents the erroneous merging of partially spliced introns into exons, improving transcript accuracy and correctly allocating reads to nascent molecules. This logic also benefits poly(A)-selected datasets by detecting and filtering internal priming artifacts^11, 12^, which can otherwise inflate false transcript assemblies. As a result, StringTie3 consistently outperforms existing methods in both short-read and long-read assembly across multiple library types, achieving higher precision and sensitivity.

**Figure 2.**
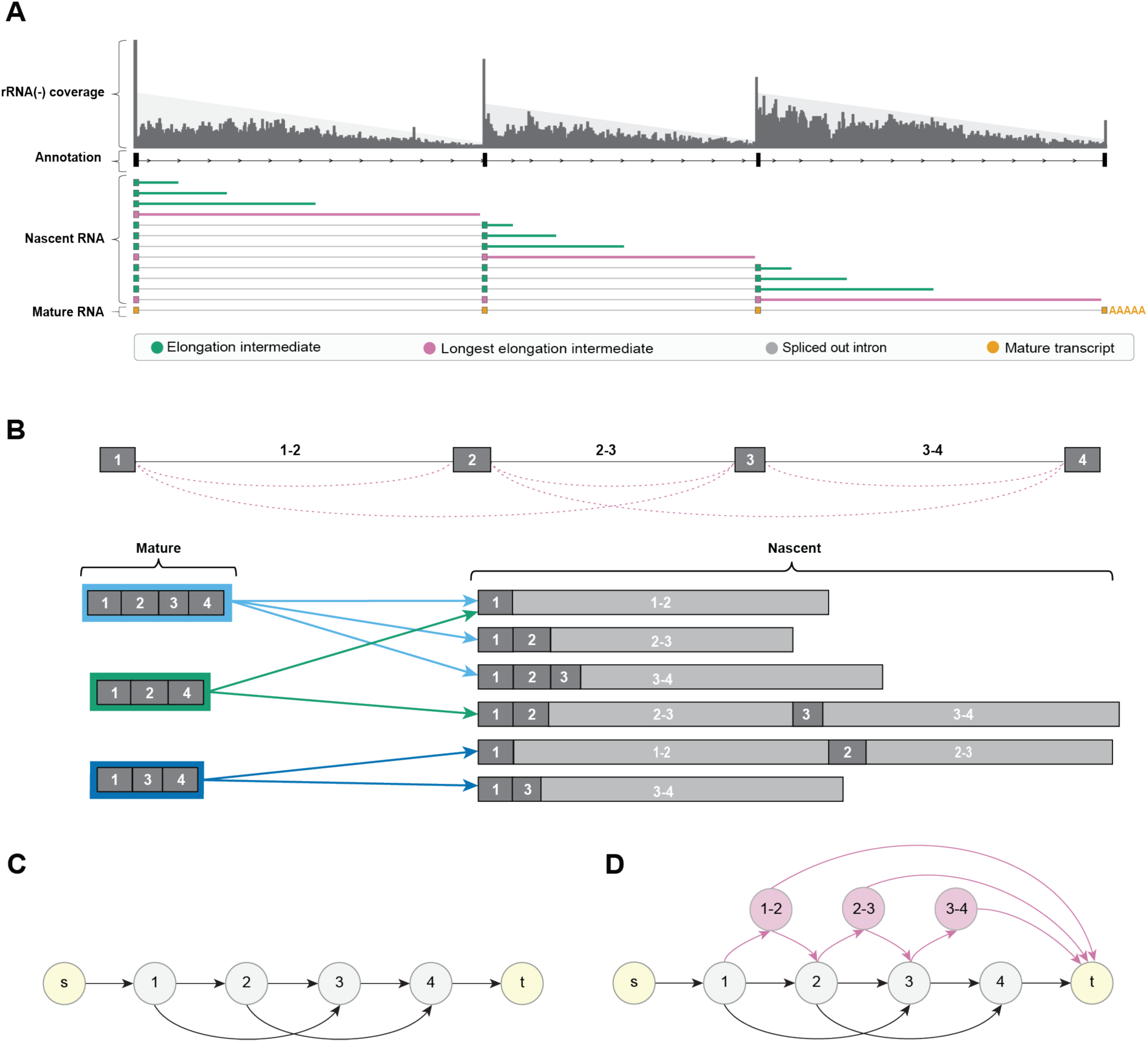
Modeling partially spliced nascent transcripts and mature isoforms within StringTie3’s splicing graph. **(A)** A hypothetical gene model illustrates how introns are progressively transcribed (green lines) and eventually spliced out (gray lines). As the RNA polymerase advances, co-transcriptional splicing removes fully transcribed introns, creating partially spliced intermediates with decreasing intron coverage. Rather than producing a separate transcript for every incremental intron extension, StringTie3 consolidates each set of co-transcriptional intermediates into a single nascent transcript (pink lines). This captures multiple maturation states for every intron. **(B)** A gene with three isoforms (all containing three exons) is transcribed into both fully spliced (mature) RNAs and partially spliced (nascent) RNAs. Although all three isoforms share the first exon, the second isoform (green) skips exon 2, and the third isoform (dark blue) skips exon 3. Each isoform produces one or more nascent intermediates; arrows indicate the longest nascent RNA for each intron. **(C)** Splicing graph constructed under the assumption that only mature isoforms are present. The two yellow nodes (s and t) mark the source and sink of the transcripts. **(D)** In nascent mode, StringTie3 adds intronic nodes and edges (shown in pink) to represent transcripts that terminate within unspliced introns. By allowing intronic nodes to connect to the sink node (t), the graph accurately models incompletely processed RNAs co-existing with mature isoforms.

By differentiating partially processed from fully mature transcripts in total RNA-seq data, StringTie3 facilitates more nuanced analyses of co- and post-transcriptional regulation, reduces reliance on specialized nascent assays, and expands the utility of rRNA-depleted libraries, which have historically been used for non-polyadenylated RNA studies (e.g., A-to-I editing, circRNAs). Ultimately, StringTie3 addresses long-standing challenges in rRNA-depleted workflows and provides an effective solution for transcriptome assembly in these datasets.

## Results

### Overview of StringTie3

To improve transcriptome assembly from rRNA-depleted (total) RNA-seq data, we developed a ‘nascent mode’ within our previously developed transcriptome assembler StringTie2^8, 14^ that explicitly accounts for partially spliced nascent transcripts (see Fig. 2 and Methods). In rRNA-depleted libraries, abundant intronic coverage often reflects transcripts that have not fully completed elongation or splicing, and failing to account for these intermediates can lead to frequent misassemblies (Fig. 1B).

Figure 2A shows how introns are progressively transcribed (green lines) and then spliced out (gray lines), generating partially spliced intermediates with decreasing intron coverage and resulting in a ‘sawtooth’ coverage pattern. Rather than building a separate transcript for each incremental nucleotide addition, StringTie3 consolidates all co-transcriptional intermediates into a single nascent transcript (pink lines), capturing every in-progress version that terminates within the same intron (Fig. 2A and 2B).

By default, StringTie assumes that most reads originate from mature transcripts (Fig. 2C). However, in rRNA-depleted data, ignoring intronic reads leads to misassemblies. To address this, nascent mode augments the splicing graph with intronic nodes and edges, allowing transcripts to terminate in unspliced introns. As shown in Fig. 2D (pink nodes and edges), this design more accurately reflects multiple elongation stages and prevents reads spanning introns from being misassembled as exons. Because splicing generally occurs co-transcriptionally, StringTie3 inserts intronic nodes between exonic nodes that are connected by a splice junction, along with edges that allow the transcript to terminate there (Fig. 2D, pink nodes and arrows). This approach ensures correct intronic termination for nascent transcripts while also enabling the accurate assembly of isoforms with retained introns, supported by reads spanning the intron and adjacent exonic nodes (see Methods).

Although the nascent mode is not expected to significantly affect the assembly of poly(A)-selected RNA-seq libraries, we observed consistently improved transcriptome reconstruction, particularly in long-read data. This improvement occurs because the nascent mode systematically removes intronic priming artifacts, which often arise from internal poly(A) tracts within introns and cause inadvertent sequencing of nascent intermediates (Supplementary Figure 1). Building on this insight, we updated StringTie3’s long-read module to detect and remove these poly(A) priming artifacts (see Methods), further improving the accuracy of long-read assemblies.

### Benchmarking on rRNA-depleted short-read datasets

To benchmark StringTie3 (nascent mode) against StringTie2 and the widely used Scallop2^15^ transcriptome assembler, we evaluated three distinct short-read, rRNA-depleted RNA-seq datasets: (1) Dorsolateral Prefrontal Cortex (DLPFC)^16, 17^, (2) Breast Cancer (tumor vs. normal pairs)^18^, and (3) Neuron Differentiation (time points from Day 0 to Day 5 of induced pluripotent stem cells)^19^. From each dataset, we randomly selected ten samples (30 samples total) and measured sensitivity and precision by comparing the assembled transcripts to a reference annotation. Because Scallop2 lacks an annotation-guided assembly mode, all three programs were run in annotation-free mode (see Methods).

As shown in Figure 3A, StringTie3 (nascent mode) outperforms StringTie2 and Scallop2 in both sensitivity and precision. Relative to Scallop2, StringTie3 achieves an average relative improvement of 2.5% and 42.0% in sensitivity and precision, respectively, across all 30 samples, and 2.9% and 13.0%, respectively, when compared to StringTie2. In 7 out of the 30 samples, Scallop2 displayed a minor (1–4%) relative gain in sensitivity over StringTie3 but generally lags by more than 40% in precision (Fig. 3A). We attribute the slight sensitivity drop in StringTie3 (vs. Scallop2) to reference transcripts sharing intron chains with assembled nascent transcripts, where coverage dips can be misconstrued as transcript boundaries (Supplementary Fig. 2). While Scallop2 and StringTie2 often interpret variable intronic coverage—common in rRNA-depleted libraries—as transcript boundaries, StringTie3 classifies these regions as nascent RNA based on surrounding intronic coverage and excludes them from the mature transcript assembly (Figure 3B, C). Notably, when such transcripts are genuinely present at coverage levels exceeding the nascent background, StringTie3 correctly assembles them as mature (Fig. 3D). Overall, these findings highlight the advantage of nascent-aware modeling for improving rRNA-depleted transcriptome assembly accuracy.

**Figure 3.**
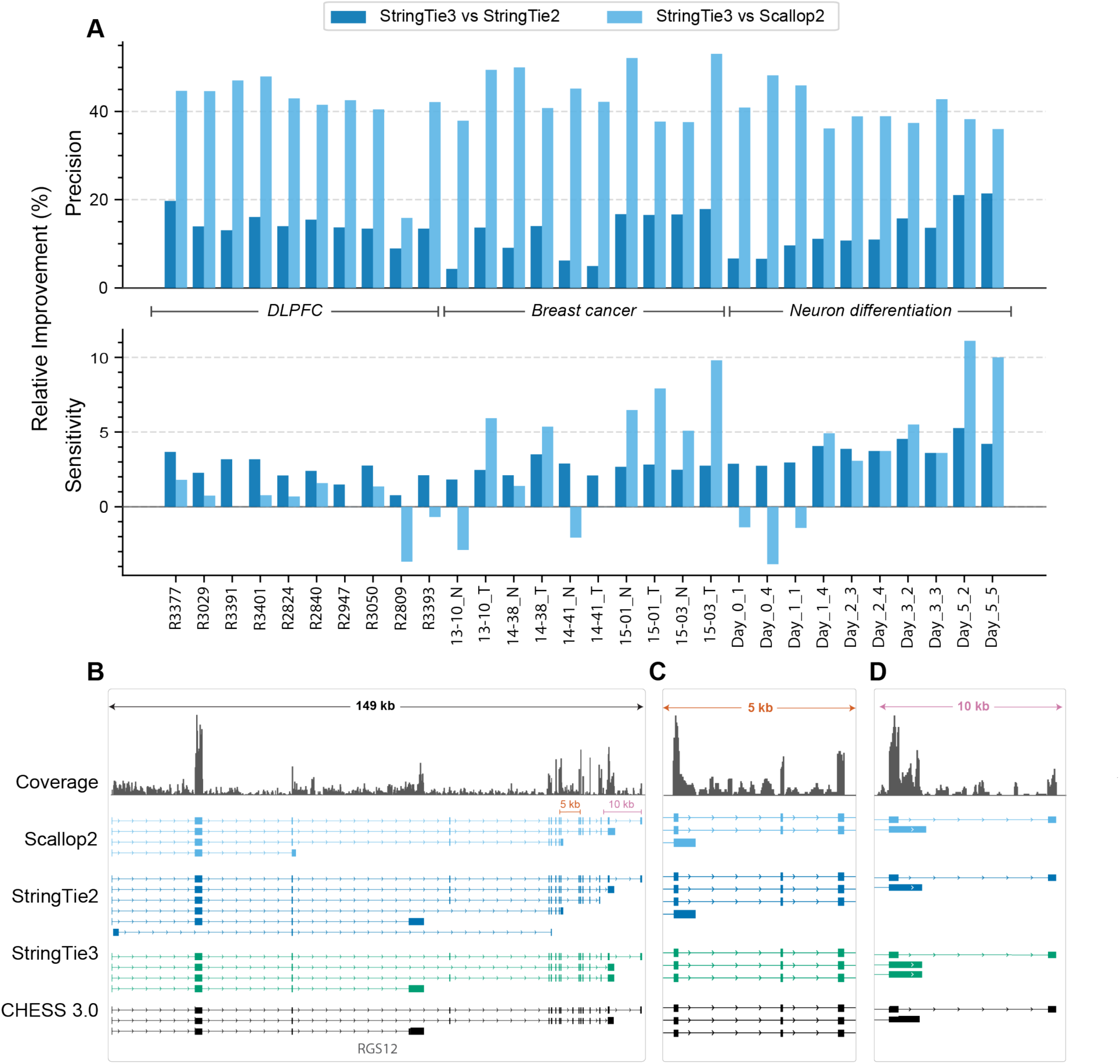
Comparison of StringTie3 (nascent mode), StringTie2, and Scallop2 in rRNA-depleted RNA-seq. **(A)** Relative changes in sensitivity (top panel) and precision (bottom panel) of StringTie3 (nascent mode) versus StringTie2 (dark blue bars) and Scallop2 (light blue bars) across 30 rRNA-depleted RNA-seq samples from three distinct datasets: Dorsolateral Prefrontal Cortex (DLPFC), Breast Cancer (tumor–normal pairs), and Neuron Differentiation (Day 0–Day 5). Positive values indicate improved sensitivity or precision for StringTie3 relative to the other assemblers, while negative values indicate a decrease. Although Scallop2 occasionally shows a 1–4% relative sensitivity increase over StringTie3, it generally lags by more than 40% in precision. **(B)** A gene locus with multiple introns and variable intronic coverage, where Scallop2 and StringTie2 interpret coverage dips as transcript boundaries (orange marker, 5 kb region), producing partial transcripts that lower precision. In contrast, StringTie3 classifies these coverage dips as nascent RNA. When the coverage drop exceeds the variability of the intronic region, StringTie3 properly assembles a transcript end that matches the reference annotation (pink marker, 10 kb region). **(C)** Zoomed view of the orange region in (B) highlighting an intronic coverage dip interpreted as a transcript end by Scallop2 and StringTie2 but not by StringTie3. **(D)** Zoomed view of the pink region in (B) (10 kb marker) showing a significant coverage drop that StringTie3 correctly treats as a transcript boundary.

### Comparison of matched poly(A)-selected and rRNA-depleted RNA-Seq datasets

To evaluate how library preparation affects transcriptome assembly, we analyzed 144 matched dorsolateral prefrontal cortex (DLPFC) RNA-seq samples, generated using either poly(A) selection or rRNA depletion. Each sample was assembled with StringTie3 (nascent mode) and StringTie2 in both annotation-guided and annotation-free modes, and we assessed precision and sensitivity against the CHESS 3.0 reference annotation^20^.

In poly(A)-selected libraries, StringTie3’s nascent mode improved precision by an average of 10.7% in annotation-guided mode and 2.9% in annotation-free mode, while maintaining essentially the same sensitivity. As expected, the gains were more pronounced in rRNA-depleted libraries, with an average precision increase of 21.4% in annotation-guided mode and 13.2% in annotation-free mode, respectively, along with an additional 2.1% rise in sensitivity for annotation-free assembly (Supplementary Table 1; see Supplementary Data 1 and 2 for per-sample statistics). Violin plots of precision and the number of reference transcripts assembled (Fig. 4A-H) confirm that StringTie3’s nascent mode consistently outperforms StringTie2 across individual samples. Additionally, for annotation-guided assemblies, the precision of rRNA(–) libraries processed with StringTie3 nearly matches that of poly(A)+ libraries, whereas rRNA(–) precision under StringTie2 remained substantially lower than that of the poly(A)+ assembly, highlighting the specific benefit of nascent mode for rRNA depleted data.

**Figure 4.**
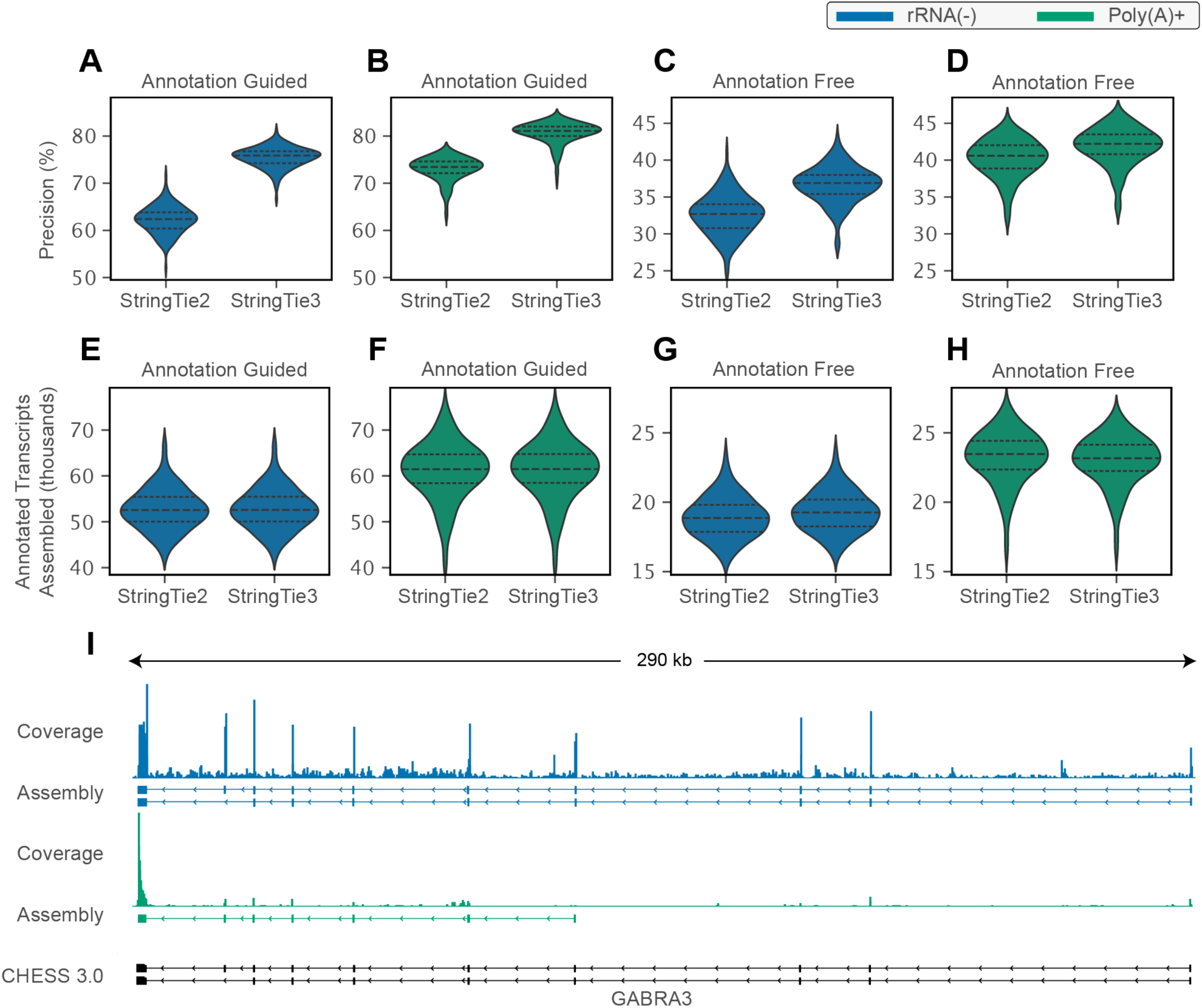
Precision, sensitivity, and coverage differences for StringTie2 vs. StringTie3 (nascent mode) in poly(A)+ and rRNA-depleted libraries. (A-D) Violin plots of precision and **(E-H)** sensitivity in poly(A)+ (green) and rRNA-depleted (blue) libraries, each evaluated in annotation-guided and annotation-free modes. Each distribution represents 144 matched DLPFC RNA-seq samples, with medians and quartiles shown by dashed lines. Across all conditions, StringTie3 (nascent mode) consistently achieves higher precision than StringTie2 while maintaining or improving sensitivity. **(I)** Coverage and StringTie3 assemblies in rRNA-depleted (blue) versus poly(A)+ (green) libraries. The rRNA-depleted data shows relatively uniform coverage across exons, enabling full-length transcript assembly, whereas coverage in the poly(A)+ library sharply declines toward the 5′ end and prevents complete assembly of the transcript. In this example, StringTie3 assembles two reference transcripts from the rRNA(-) library, while in the poly(A)+ library, low 5′ coverage yields a single truncated transcript.

We observed that poly(A)-selected libraries generally exhibit higher sensitivity, as they primarily capture mature transcripts, whereas rRNA-depleted datasets also include incomplete nascent molecules. However, most reference transcripts detected exclusively in poly(A)+ assemblies but absent from rRNA(–) libraries have low TPM values (Supplementary Fig. 3A). Reference transcripts found only in rRNA-depleted libraries tend to be longer, suggesting that poly(A)+ selection, particularly in lower-quality samples (e.g., biopsies), biases transcript detection toward shorter isoforms due to 5′ degradation (Supplementary Fig. 3B). By comparison, rRNA depletion captures transcripts more evenly along their entire length, yielding a broader spectrum of RNA isoforms^21^. These length distributions, combined with nascent-mode assembly, indicate that rRNA depletion recovers isoforms that would otherwise be missed by poly(A)+ selection alone (Fig. 4I). Consequently, an rRNA-depleted, nascent-aware approach offers a more comprehensive view of the transcriptome, including longer isoforms that might be missed by poly(A)-selected protocols.

Both rRNA-depleted and poly(A)-selected library assemblies showed improved accuracy relative to the reference annotation. Because each sample included paired rRNA(–) and poly(A)+ libraries, we assessed whether StringTie3’s nascent mode would increase the overlap between these two assemblies compared to StringTie2. In annotation-guided mode, the average number of assembled transcripts unique to rRNA(–) decreased by 38%, and those unique to poly(A)+ decreased by 20%, with little change in their shared transcript count (Supplementary Fig. 4). In annotation-free mode, the count of transcripts exclusive to rRNA(–) decreased by 7%, and those unique to poly(A)+ dropped by 13%, again with minimal impact on the overlap. Per-sample counts illustrating these comparisons are provided in Supplementary Data 3. Overall, these results indicate that explicitly modeling co-transcriptional splicing in nascent mode improves assembly accuracy relative to the reference annotation and brings rRNA(–) and poly(A)+ assemblies into closer agreement for matched samples.

### Long-read and hybrid assembly performance

We next evaluated StringTie3 on a hybrid dataset of short- and long-read RNA-seq from induced pluripotent stem cells (iPSCs) differentiating into neurons over five days. The dataset included rRNA-depleted short-read and poly(A)+ long-read sequencing at days 0, 3, and 5.

### Long-read assembly

We evaluated StringTie3 in long-read mode and compared it to StringTie2 across nine samples from the iPSC-to-neuron differentiation time course. Unlike StringTie2, StringTie3’s long-read module discards long reads terminating with a poly(A) tract aligned to the genome—likely reflecting incomplete nascent transcripts arising from poly(A) priming artifacts (Supplementary Fig. 5). We ran StringTie3 in both default and nascent modes. In long-read mode, StringTie’s default parameters (for both versions) are highly permissive, assembling any transcript supported by a single long read.

Using default parameters in annotation-free mode (averaged over the nine samples), StringTie3 nascent mode improved precision by 41.7% relative to StringTie2, with only a minor −1.8% drop in sensitivity (Supplementary Fig. 6C and 6G). Nascent mode provided only a slight additional improvement in annotation-free conditions, as expected given that the long reads were poly(A)-selected (Fig. 5A). When sensitivity was matched, StringTie3 consistently outperformed StringTie2 in precision. Under annotation-guided mode, StringTie3 nascent mode increased sensitivity by 40% and precision by 76%, substantially enhancing isoform reconstruction (Fig. 5B, and Supplementary Fig. 6A and 6E). Notably, violin plots (Supplementary Fig. 6A) show that StringTie3 produces a tighter precision range, with some samples far exceeding the mean improvement.

**Figure 5.**
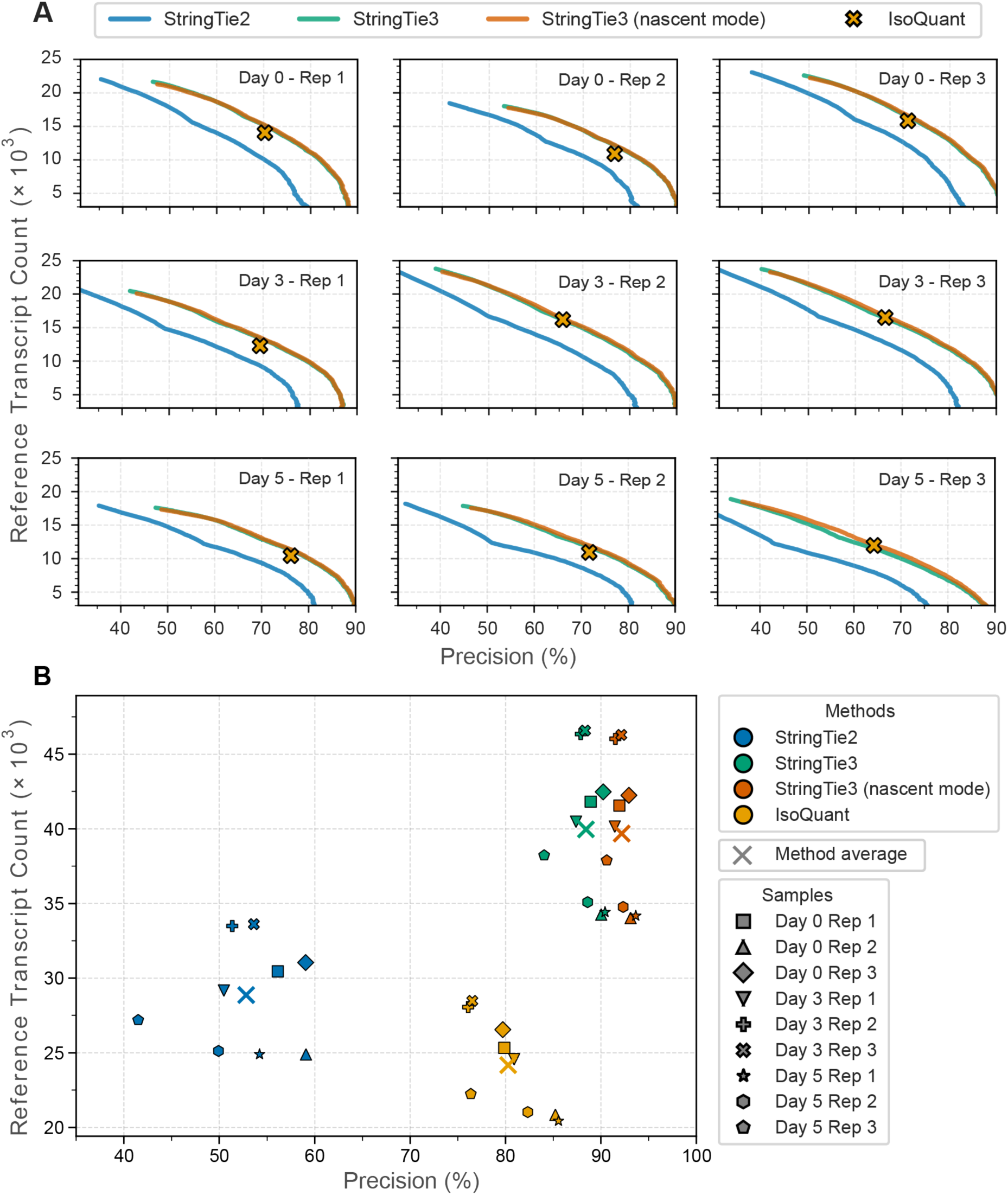
Long-Read Assembly Performance of StringTie3, StringTie2, and IsoQuant. **(A)** Annotation-free mode. Each panel plots precision (x-axis) vs. number of matched reference transcripts (y-axis) for a single sample at days 0, 3, and 5 (three replicates each). StringTie2 is shown in blue, StringTie3 in green, StringTie3 nascent mode in orange, and IsoQuant in yellow. Curves are generated by varying the minimum coverage threshold used to filter low-coverage transcripts before calculating sensitivity and precision. On average, StringTie3 achieves higher precision than StringTie2, and nascent mode adds only a minor improvement in annotation-free conditions. **(B)** Annotation-guided mode. Scatter plots of precision vs. matched reference transcript counts for the same nine samples. StringTie3 shows substantial gains in both precision and sensitivity over StringTie2 and IsoQuant.

We next benchmarked StringTie3 against IsoQuant^22^, recently identified as the most accurate long-read assembler in a benchmarking study by Su et al.^23^. As shown in Figure 5, StringTie3 outperforms IsoQuant under both annotation-free (Fig. 5A) and annotation-guided (Fig. 5B) modes. In annotation-free mode, at equivalent sensitivity, StringTie3 consistently achieved higher precision in six of the nine samples and was comparable in the remaining three. Under annotation-guided conditions, StringTie3 boosted precision by 15% relative to IsoQuant and identified 64% more reference transcripts.

### Hybrid assembly

Because StringTie uniquely supports hybrid assembly from short- and long-read data, we compared StringTie3 only to StringTie2 for this dataset. Hybrid data combine the high coverage of short reads with the full-length isoform resolution of long reads, facilitating improved detection of complex splicing events ^14^. Given that the short-read sequencing is total RNA-seq, StringTie3’s nascent mode is particularly well suited to model co-transcriptional splicing in this dataset.

In annotation-free mode, StringTie3 increased precision by 53.5% on average, albeit with a 4.1% decrease in sensitivity (Supplementary Figure 6D and 6H). This reduction largely results from more stringent filtering of partial or low-coverage transcripts in hybrid-mode—reads that might otherwise be allocated to mature isoforms are recognized as nascent, reducing apparent coverage below the default detection threshold. However, in annotation-guided mode, StringTie3 demonstrates a 53.9% relative gain in precision along with a 28.8% gain in sensitivity, indicating that co-transcriptional modeling and poly(A) artifact filtering work synergistically to enhance transcript reconstruction in hybrid RNA-seq datasets (Supplementary Figure 6B and 6F).

Moreover, while the average improvement is substantial, examining the violin-plot distributions (Supplementary Fig. 6B) shows a tighter precision range for StringTie3, with certain samples far exceeding the mean. In one example, precision nearly doubled from 47.6% (StringTie2) to 83.4% (StringTie3 with nascent mode).

### Disentangling nascent and mature RNA to probe transcriptional and post-transcriptional regulation

One important application of StringTie3’s ability to quantify both nascent and mature RNA species from rRNA-depleted total RNA-seq data is investigating the regulatory dynamics of gene expression. To demonstrate this, we analyzed RNA-seq data from two biologically relevant contexts where both transcriptional and post-transcriptional regulation are at play: (1) Argonaute knockouts in HCT116 cells and (2) tumor–normal comparisons in breast cancer samples. These case studies demonstrate how quantifying nascent and mature RNA reveals whether changes in gene expression are primarily driven by transcription or shaped by post-transcriptional mechanisms.

In all subsequent experiments, we used StringTie3 in nascent mode to identify and quantify transcripts and performed differential expression analyses for nascent and mature RNA components. To minimize potential noise from incorrectly predicted novel transcripts, we restricted our analysis to transcripts present in the reference annotation and their nascent counterparts (see Methods).

### Argonaute (AGO1/AGO2) knockouts reveal post-transcriptional buffering

To demonstrate StringTie3’s ability to distinguish nascent from mature RNA, we analyzed data from Chu et al. (2020) in which AGO1 and AGO2 were knocked out individually or in combination (AGO1/2 and AGO1/2/3) in HCT116 colorectal cancer cells, followed by rRNA-depleted RNA-seq. Argonaute proteins are core components of RNA interference (RNAi) and microRNA pathways and have also been implicated in transcriptional regulation^24–26^.

#### Minimal mature-RNA changes in single knockouts

To assess how Argonaute proteins influence transcription and mRNA abundance, we compared nascent and mature RNA expression across various knockout conditions. Figure 6 (A-D) plots nascent (x-axis) vs. mature (y-axis) log₂ fold changes for each knockout relative to wild-type. In single knockouts (Fig. 6A, B), many genes show large changes in nascent RNA without corresponding changes in mature RNA, suggesting that although Argonaute proteins modulate transcription, other family members buffer the final mRNA levels. Conversely, removing both AGO1 and AGO2 or all three (Fig. 6C, D) eliminates this buffering capacity and induces differential expression in both nascent and mature fractions. These findings align with earlier reports of Argonaute-mediated transcriptional regulation^24–26^.

**Figure 6.**
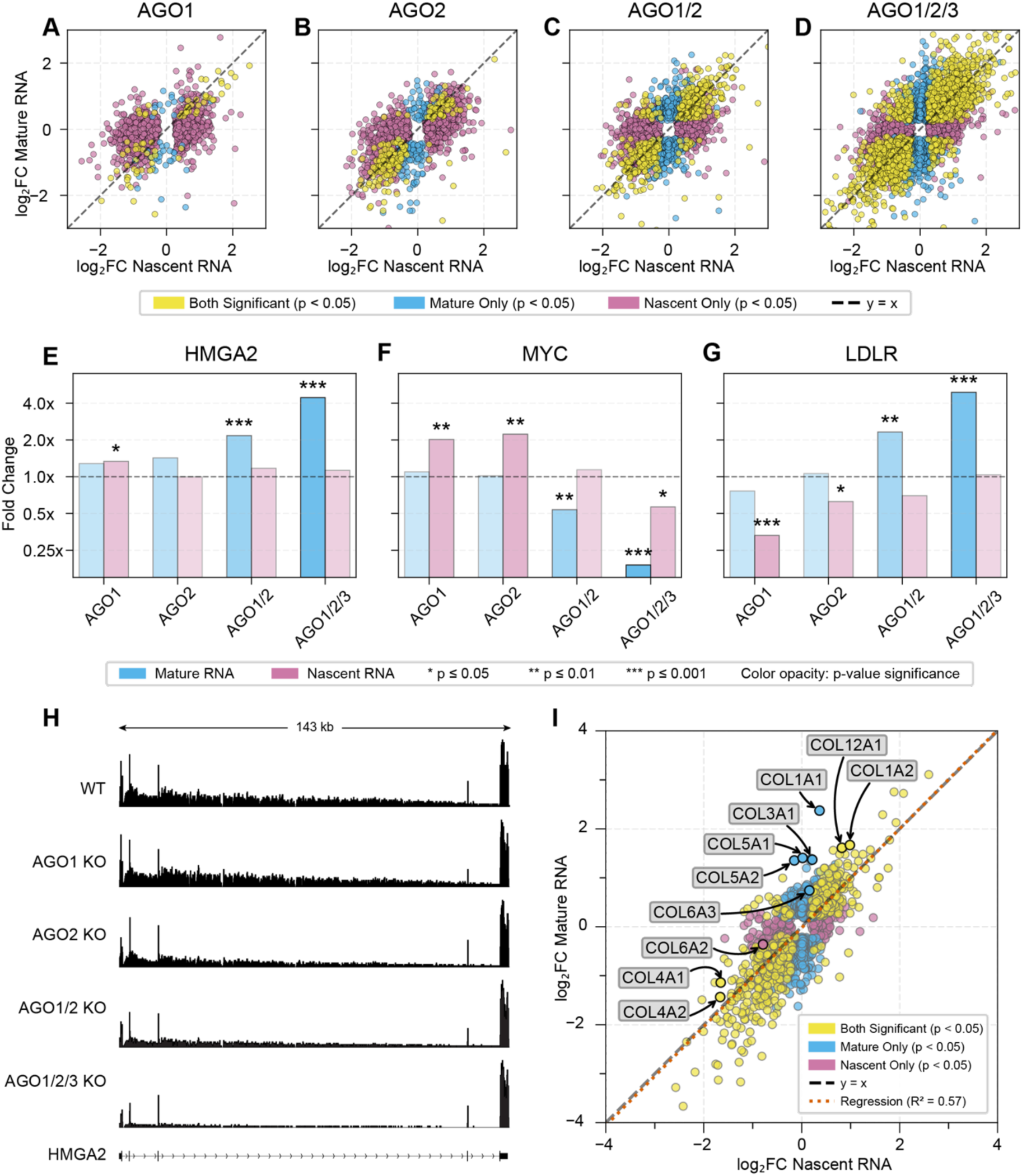
Argonaute Knockouts Reveal Transcriptional vs. Post-Transcriptional Regulation (A–D) Scatter plots of nascent (x-axis) vs. mature (y-axis) log₂ fold changes for AGO1 *(A)*, AGO2 *(B)*, AGO1/2 *(C)*, and AGO1/2/3 *(D)* knockouts relative to wild type. Dots are colored yellow if FDR-adjusted p < 0.05 in both analyses, blue if significant only in mature, and reddish purple if significant only in nascent**. (E–G)** Fold-change bar plots for HMGA2, c-MYC, and LDLR, each comparing nascent vs. mature fold changes for single (AGO1, AGO2) and double/triple (AGO1/2, AGO1/2/3) knockouts. **(H)** Coverage tracks of HMGA2 in WT, AGO1 KO, AGO2 KO, AGO1/2 KO, and AGO1/2/3 KO cells, illustrating how multi-Argonaute depletion shifts the mature-to-nascent RNA ratio, consistent with panel E. **(I)** A scatter plot of nascent (x-axis) vs. mature (y-axis) log₂ fold changes—mirroring panels (A–D) but applied to 21 matched tumor–normal rRNA-depleted breast cancer samples. Notably, collagens appear above the diagonal (y = x), indicating stronger post-transcriptional upregulation than co-transcriptional changes.

#### HMGA2: a post-transcriptional target of let-7

HMGA2 is an oncogene regulated post-transcriptionally by the let-7 microRNA family^27^. In single AGO1 or AGO2 knockouts, HMGA2 mRNA levels remain unchanged, whereas double AGO1/2 depletion leads to a two- and four-fold increase (Fig. 6E, H). Nascent HMGA2 expression remains stable across conditions, suggesting that partial Argonaute loss has little effect on transcription. The lack of change in single knockouts indicates that remaining Argonautes buffer mRNA levels, while combined Argonaute depletion (AGO1/2 and AGO1/2/3) leads to significant accumulation of HMGA2–highlighting the role of miRNA-directed decay in maintaining normal HMGA2 expression.

#### c-MYC: multiple layers of AGO-mediated regulation

c-MYC is an oncogene frequently overexpressed in human cancers and is subject to complex post-transcriptional regulation^28^. In single AGO1 or AGO2 knockouts, nascent c-MYC levels double while mature RNA levels remain steady (Fig. 6F), suggesting that each Argonaute represses c-MYC transcription, and the resulting mRNA abundance is buffered post-transcriptionally by the remaining Argonautes. This observation is consistent with prior evidence that AGO1 can repress MYC transcription^24–26^. In AGO1/2 double knockouts, mature c-MYC decreases by ∼50% despite stable nascent levels, and triple AGO1/2/3 knockout reduces both fractions. The decrease in mature c-MYC in the double and triple knockouts mirrors results from Chu et al.^29^, who found that Argonaute binding in the c-MYC 3′UTR does not reliably predict repression. By separately measuring nascent and mature RNA, we show that partial Argonaute loss primarily affects transcription, whereas multi-Argonaute depletion reduces mature c-MYC abundance at both the transcriptional and post-transcriptional levels.

#### LDLR: distinguishing transcriptional vs. post-transcriptional effects

LDLR is a key regulator of cholesterol homeostasis that is subject to microRNA-mediated transcriptional and post-transcriptional regulation^30, 31^. In single AGO1 or AGO2 knockouts, nascent LDLR levels decrease, yet mature LDLR remains unchanged (Fig. 6G), suggesting that Argonaute-mediated transcriptional changes are buffered at the mRNA level by the remaining AGOs. However, in double or triple knockouts, mature LDLR levels increase two- to four-fold despite minimal changes in nascent RNA, revealing a dominant post-transcriptional effect. These observations suggest that LDLR mRNA abundance relies primarily on microRNA-mediated post-transcriptional silencing: losing a single Argonaute reduces transcription but does not alter mature LDLR abundance, while depleting multiple Argonautes disrupts post-transcriptional repression and leads to the observed increase in mature LDLR.

#### Regulatory insights from StringTie3’s nascent and mature RNA analysis

Across HMGA2, c-MYC, and LDLR, StringTie3’s nascent-mode analysis highlights cases where nascent RNA levels diverge from mature RNA, indicating transcriptional and post-transcriptional regulation. For instance, in c-MYC and LDLR, single Argonaute knockouts alter nascent levels with minimal effect on mature RNA, reflecting redundant buffering by remaining Argonautes. Conversely, removing two or more Argonautes disrupts both nascent and mature RNA abundance, demonstrating the collective role of Argonautes in stabilizing steady-state mRNA levels. This observation is consistent with previous findings that Argonautes act redundantly to buffer gene expression fluctuations, even when one family member is lost^32^. See Supplementary Data 4–7 for a detailed overview of these differential expression analyses. By analyzing nascent vs. mature fractions, StringTie3 enables researchers to determine whether expression changes stem from altered transcription, post-transcriptional buffering, or both— mechanisms that are often obscured in poly(A)-selected or mature-only RNA-seq.

#### Application of StringTie3 to breast cancer data

Building on the Argonaute findings, we next applied StringTie3 to an rRNA-depleted dataset of 21 matched tumor–normal breast cancer samples^18^. While the Argonaute experiments illustrate how nascent-mode quantification disentangles transcriptional from post-transcriptional effects in a controlled knockout model, the breast cancer dataset shows that the same approach can provide insights into complex regulatory mechanisms in a disease context.

#### Post-transcriptional regulation of collagens in breast cancer

Collagens are increasingly recognized as key factors in tumor dormancy, immune evasion, and treatment response^33^. In our nascent vs. mature differential expression analysis (Methods), all ten differentially expressed collagen genes lie above the x = y line in a nascent vs. mature log₂ fold-change plot (Fig. 6I), indicating stronger upregulation in their mature fractions than in nascent RNA. This trend suggests a shared post-transcriptional mechanism affecting the collagen family. For example, COL5A1 showed negligible changes at the nascent level but was over two-fold higher in its mature fraction, consistent with recent findings linking its overexpression to breast cancer progression and chemoresistance^34^. Similarly, COL3A1 undergoes only minor changes in nascent RNA but is substantially elevated in its mature form, aligning with earlier studies that report COL3A1 stabilization in breast cancer^35^.

Several pathways may explain the discrepancy between nascent and mature collagen expression. miR-29, a known tumor suppressor microRNA, negatively regulates collagen^36^, and its downregulation or loss (e.g., miR-29b/c) has been associated with elevated expression of multiple collagen genes^37^. Additional factors, such as m^6^A modifications^35, 37^ or lncRNA “sponges” that sequester collagen-targeting miRNAs^38^, may also contribute to the pronounced differences we observe between nascent and mature collagen RNA. By disentangling these two RNA pools, StringTie3 highlights a post-transcriptional signature underlying collagen overexpression in breast tumors.

#### Other genes with nascent–mature discrepancies

Beyond collagens, we identified numerous genes whose nascent and mature expression levels diverge. For instance, FN1 (fibronectin), an extracellular matrix protein that interacts with collagen, increased by over four-fold at the nascent level but more than six-fold in mature RNA, consistent with reports that miR-29 loss can jointly upregulate FN1 and collagens^37^. In contrast, the tumor suppressors FHL1 and TNS1 declined by approximately three- and five-fold, respectively, in nascent RNA, yet dropped by more than eight- and twelve-fold in their mature fraction. This pattern suggests additional post-transcriptional regulation beyond the observed transcriptional changes. For instance, FHL1 is repressed by miR-410^39^ while TNS1 is targeted by miR-942^40^. Other genes, including CRTAP, ALAD, and RGS5, showed elevated nascent RNA but reduced mature levels, indicating that post-transcriptional mechanisms can counterbalance transcriptional upregulation. Refer to Supplementary Data 8 for the complete differential expression analysis in this breast cancer cohort.

## Discussion

Accurately assembling rRNA-depleted (total) RNA-seq data requires disentangling nascent transcripts from fully processed mature isoforms, a challenge that standard assemblers often overlook. Here, we introduced StringTie3, an updated version of the widely used StringTie algorithm, specifically engineered to address these complexities. By modeling co-transcriptional splicing and filtering internal poly(A)-priming artifacts, StringTie3 substantially improves transcript assembly across a variety of library types—short-read, long-read, and hybrid RNA-seq datasets.

Our short-read benchmarks show that StringTie3 (nascent mode) consistently outperforms existing tools (StringTie2, Scallop2) in rRNA-depleted datasets from multiple tissue and cell types. Achieving over 20% and 50% higher precision relative to StringTie2 and Scallop2, respectively—while maintaining or improving sensitivity—highlights how co-transcriptional modeling prevents the high intronic coverage typical of rRNA-depleted libraries from introducing assembly noise. Moreover, analysis of matched rRNA-depleted and poly(A)-selected libraries shows that nascent mode aligns these datasets more closely, underscoring its potential to unify transcript sets across diverse sample-preparation methods.

In long-read applications, StringTie3’s refined module discards incomplete transcripts arising from intronic poly(A)-priming artifacts, substantially improving assembly. In annotation-guided mode, precision increases by up to 75% and sensitivity by 37% compared to StringTie2. In annotation-free mode, StringTie3 surpasses both StringTie2 and IsoQuant at the same sensitivity level, achieving higher precision and a closer match to the reference annotation. Hybrid assemblies also benefit from nascent mode, especially in datasets that pair rRNA-depleted short-read coverage with the isoform resolution of poly(A)-selected long reads, thereby enabling more accurate reconstruction of complex splicing events.

Argonaute knockout experiments illustrate how StringTie3’s nascent-mode quantification can distinguish transcriptional changes from post-transcriptional effects. Single Argonaute knockouts elicit large shifts in nascent RNA but little alteration of mature RNA, whereas double or triple knockouts disrupt both fractions. Such findings illustrate the collective role of Argonautes in maintaining steady-state mRNA levels. Similarly, in breast cancer samples, nascent-mode analysis reveals a unifying “collagen signature,” in which differentially expressed collagen genes show greater upregulation in their mature fractions compared to nascent RNA as well as revealing other pathways governed by a combination of transcriptional and post-transcriptional regulation—insights typically obscured when only mature RNA is profiled.

StringTie3’s nascent flag streamlines nascent transcription profiling without requiring specialized protocols such as GRO-seq, PRO-seq, or 4sU-seq. The resulting assemblies capture incomplete nascent RNAs alongside mature transcripts in the same rRNA-depleted sample. In doing so, StringTie3 broadens the applicability of total RNA-seq and opens new avenues to investigate transcriptional and post-transcriptional dynamics. Clinical samples often cannot accommodate nascent RNA assays, which require actively transcribing cells, so StringTie3 offers a practical solution for characterizing both nascent and mature transcripts simultaneously. Moreover, poly(A) selection in degraded or biopsy-derived samples often biases analyses toward shorter isoforms, whereas combining rRNA libraries with StringTie3’s improved assembly better captures full-length transcripts, expanding the scope of total RNA-seq in translational research.

By explicitly modeling co-transcriptional splicing and filtering spurious poly(A)-priming events, StringTie3 fully leverages total RNA-seq data: reducing assembly errors, boosting precision and sensitivity, and enabling the separation of nascent from mature RNAs in one experiment. This comprehensive approach not only illuminates hidden layers of RNA regulation but also lays the groundwork for a more complete exploration of transcriptomes, particularly in scenarios where poly(A)-selected libraries or specialized nascent-capture methods are impractical.

## Methods

### StringTie3 Algorithm

StringTie3 extends the original StringTie algorithm^8^ to improve transcriptome assembly, particularly from rRNA-depleted (total) RNA-seq data, through two main innovations: a "nascent mode" for modeling co-transcriptional splicing and a refined long-read module for filtering artifacts.

Similar to the original StringTie algorithm, StringTie3 reconstructs transcripts at each gene locus by first building a splice graph from overlapping read alignments in that locus. The nodes in this splice graph represent contiguous genomic regions and edges represent reads—either single reads or paired-end fragments (hereafter referred to as *transfrags*)—that align across two such nodes in the correct 5′ to 3′ orientation. It is important to note that nodes do not necessarily correspond to complete exons but may represent partial exons, as illustrated by node 3 in Figure 7A. Two additional nodes, a global source (*s*) and a sink (*t*), are added to the graph so that any path from source to sink represents a potential transcript.

**Figure 7.**
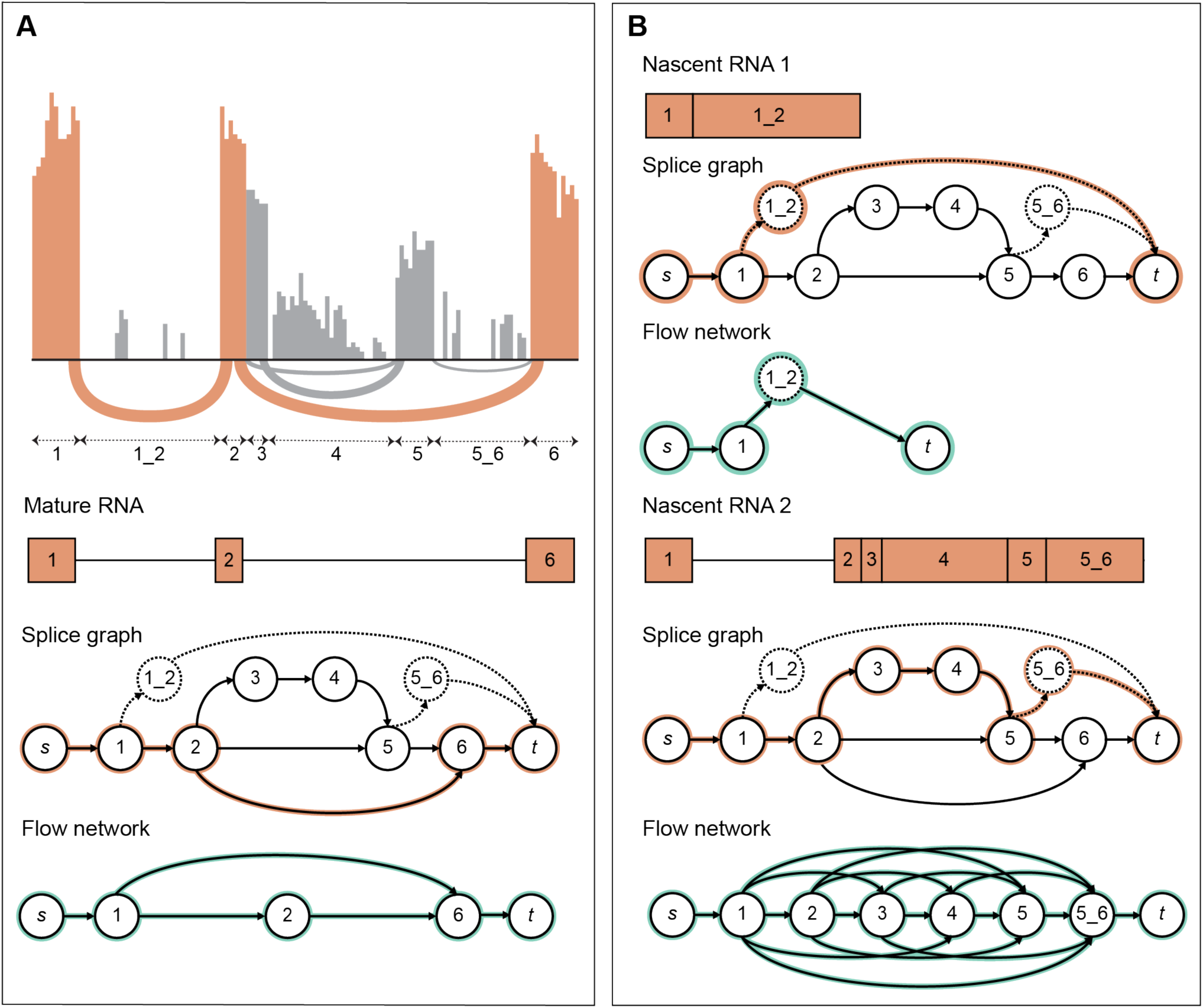
StringTie3’s nascent mode algorithm. **A**. Candidate transcript selection and quantification in a gene locus. Read coverage is shown at the top, with regions corresponding to the candidate transcript highlighted in orange. Arcs below the coverage plot represent splice junctions supported by spliced reads; thicker arcs indicate stronger read support. The splice graph, with the candidate transcript (i.e., the heaviest path) highlighted in orange, is shown below. Arrows beneath the coverage plot indicate the genomic regions corresponding to the nodes in the splice graph. Dashed nodes and edges (e.g., nodes 1_2 and 5_6) represent intronic nodes that, due to sparse coverage, are not included in the splice graph when using the non-nascent mode. A flow network (highlighted in green) is then constructed using all nodes from the heaviest path, with edges connecting two nodes if a transfrag starts at one and ends at the other. **B**. Nascent transcript quantification. Splice graphs and flow networks corresponding to the two nascent transcripts derived from the transcript in panel A are shown. The paths of the nascent transcripts are highlighted in orange in the splice graphs.

To identify transcripts at a locus, StringTie3 iteratively selects a candidate transcript as the heaviest path from source to sink (i.e., the path with the highest overall read coverage) and constructs a generalized flow network restricted to the nodes along that path. Formally, the generalized flow network is defined as a quadruple 𝑁 = (𝐺, 𝑠, 𝑡, 𝑐), where 𝐺 = (𝑉, 𝐸) is a directed graph with 𝑉 representing the nodes along the candidate transcript path, 𝐸 is the set of directed edges connecting two nodes if a transfrag starts at one and ends at the other, 𝑠 ∈ 𝑉 and 𝑡 ∈ 𝑉 are the source and sink nodes of the network, and 𝑐: 𝐸 → ℛ^+^ is a function assigning to each edge a capacity equal to the number of transfrags supporting it. A function 𝑓: 𝐸 → ℛ^+^ is a *flow* over the network *N* if the following two conditions are satisfied:

1. 0 ≤ 𝑓(𝑒) ≤ 𝑐(𝑒), for every 𝑒 ∈ 𝐸,
2. ∑_(*u,v*)∈*E* (_ 𝛾(𝑣) 𝑓:(𝑢, 𝑣)< = ∑_(*v*,*w*)∈(_ 𝑓((𝑣, 𝑤)), for every node 𝑣 ∈ 𝑉, 𝑣 ≠ 𝑠, 𝑡.

where 𝛾(𝑣) is a gain factor associated with each node 𝑣 that accounts for coverage bias in RNA-seq data. This factor is computed during splice graph construction as:

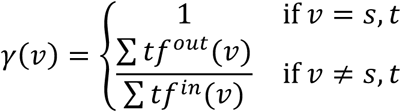

where 𝑡𝑓*^in^*(𝑣) and 𝑡𝑓*^out^* (𝑣) represent the sets of transfrags entering and exiting node 𝑣, respectively. The value of the flow is defined as |𝑓| = ∑*_(s,v_*_)∈*E*_ 𝑓(𝑠, 𝑣).

Once the flow network is built, StringTie3 estimates the abundance of the candidate transcript by solving a *maximum flow problem* that identifies a flow *f* with maximum value in the network *N*. This flow value is then used to compute the transcript’s abundance by attributing a corresponding fraction of the coverage from all nodes along the path to the transcript, reflecting the proportion of transfrags explained by this transcript relative to the total coverage. The intuition behind this flow network design is that StringTie3 attempts to stitch together compatible fragments that collectively account for the maximum number of reads supporting the underlying transcript path. In default mode (non-nascent), after the transfrags that contribute to the maximum flow are removed from the splice graph the process repeats until no remaining path meets a minimum coverage threshold.

To address the abundance of incomplete, nascent transcripts common in total RNA-seq, StringTie3 incorporates a ’nascent mode’ that explicitly models these co-transcriptional splicing intermediates, distinguishing them from fully processed, mature isoforms. This mode is activated using either the `--nasc` or `-N` flag. The `--nasc` flag includes the assembled nascent intermediates (representing partially spliced forms) in the output GTF file, whereas the `-N` flag performs the internal modeling and read assignment adjustments but excludes these nascent transcript models from the final output, reporting only mature forms. The key algorithmic modifications enabling this are:

#### 1. Intronic Node Representation

In nascent mode, StringTie3 systematically introduces ’intronic nodes’ into the splice graph at every exon-exon splice junction. An edge is added connecting each intronic node to the terminus (sink node). This structure explicitly represents unspliced introns and allows transcript paths to terminate within them, modeling incomplete nascent RNA molecules.

This approach differs from the original StringTie algorithm, which typically incorporates intronic regions only when there is direct read evidence supporting their inclusion—such as reads spanning intron–exon boundaries. In contrast, StringTie3 includes intronic nodes even in the absence of such spanning reads (e.g., node 1_2 in Figure 7A), as long as the intron shows non-zero coverage. The original method is sensitive to the variable and often low read coverage found in introns, frequently misinterpreting coverage gaps as transcript start or end points. This can result in inappropriate edges from the source or to the sink node, prematurely terminating transcript paths within introns or generating fragmented assemblies based on patchy coverage.

To address this issue, StringTie3 creates additional intronic nodes to ensure that the entire intron is represented, even when nodes from the original algorithm cover only portions of the intronic region. Formally, in nascent mode, if 𝑛*_i_*, and 𝑛*_i_*_+1_are two nodes in the original splice graph that are linked by a splice junction, and the intron between them has non-zero coverage, then StringTie3 ensures there is a path 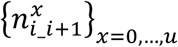 in the splice graph that spans the entire intronic region. In this path, 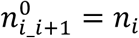 and 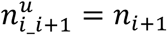, and each pair of consecutive nodes 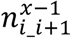 and 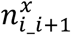, for 𝑥 = 1, …, 𝑢, are adjacent in the genomic coordinate space. New nodes are added as needed to represent regions of the intron not already covered by nodes in the original graph. If no edge exists between any pair 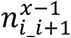 and 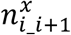, for 𝑥 = 1, …, 𝑢 − 1, StringTie3 adds a virtual transfrag with a minimal abundance 𝜃 (default: 1). The final intronic node 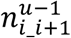 is also connected to the sink node by another virtual transfrag with abundance 𝜃, explicitly modeling nascent transcripts.

For example, Figure 7A illustrates a case where the original graph includes nodes 3, 4, and 5 within the intron between nodes 2 and 6. StringTie3 adds a new node, 5_6, to span a previously unrepresented region and connects it to both node 5 and the sink. This approach enables StringTie3 to allocate intronic reads to co-transcriptional intermediates and avoid misassemblies caused by uneven coverage. To achieve this, it uses the maximum flow algorithm described above to estimate the proportion of reads attributable to intronic nodes, distinguishing between alternative isoforms and nascent transcription.

#### 2. Heaviest Path & Flow Update

The core iterative process of identifying the heaviest path (representing the most abundant mature isoform) and assigning coverage using the maximum-flow algorithm remains similar to that of the original StringTie. However, after identifying a candidate transcript as the heaviest path in the splice graph and computing its abundance using the maximum-flow algorithm, StringTie3 removes all transfrags that contributed to that transcript and proceeds to identify all nascent transcripts associated with it. Coverage is then assigned to each nascent path by constructing a separate flow network for each one (see Figure 7B). Once the abundance of a nascent transcript is computed, the transfrags associated with it are removed. Only after all nascent transcript abundances have been estimated does StringTie3 search for the next heaviest path in the updated graph.

StringTie3 refines the long-read module introduced in StringTie2^9^ by adding two complementary filters that reduce spurious isoforms caused by intronic poly(A) priming. For single-exon alignment, the algorithm scans the first and last 25 bp that map to the genome. Reads in which at least 80% of the aligned 3’ bases are adenines (or 5’ bases are thymines), or that contain a homopolymer run of more than 10 consecutive A/T bases, are flagged as poly(A)-priming artifact and discarded. When the nascent mode is enabled, the remaining long-read assemblies are evaluated locus by locus: a candidate isoform is labelled a nascent intermediate when its exon chain matches the 5′ portion of a longer transcript yet terminates within one of that transcript’s introns, consistent with co-transcriptional splicing in which the final intron has not yet been removed. Whether these intermediates appear in the final GTF is user-controlled: `-- nasc` retains them and tags each record as “nascent,” whereas `-N` performs the same classification but omits nascent models from the output file.

### Reference Genome and Annotation

All RNA-seq datasets were aligned to the human reference genome GRCh38 (RefSeq accession GCF_000001405.39), excluding pseudoautosomal regions on chromosome Y and alternative scaffolds. Guided transcript assembly and accuracy assessment were performed using the CHESS version 3.0.1 annotation, filtered to retain only full-length protein-coding and long non-coding RNA (lncRNA) transcripts. The filtered annotation is available at ftp://ftp.ccb.jhu.edu/pub/StringTie3.

### Alignment

#### Short-Read Data

Short reads were aligned using HISAT2 (v2.2.1)^41^. A genome index was built incorporating splice sites and exons from the CHESS 3.0.1 annotation using `hisat2-build -p 16 -- exon genome.exon --ss genome.ss genome.fa hisat_index`. Alignments were performed with `hisat2 -p 8 -x hisat_index -1 R1.fastq -2 R2.fastq -S aligned.sam.` For strand-specific libraries (fr-firstrand), the --rna-strandedness RF flag was added.

#### Long-Read Data

Long reads (ONT) from the neuron differentiation dataset were aligned using minimap2 (v2.28- r1209)^42^. A genome index was built using `minimap2 -k14 -d mm2_index.mmi`. Alignments were performed with minimap2 -ax splice --junc-bed short_read_junctions.bed mm2_index.mmi sample.fastq > aligned.sam. The -k14 parameter was used as recommended for ONT reads. The short_read_junctions.bed file contained splice junctions identified from the corresponding short-read samples after alignment and filtering (see below).

#### Alignment File Processing

Resulting SAM files were converted to sorted, indexed BAM format using samtools (v1.20). To remove spurious alignments potentially arising from library preparation artifacts or mapping errors, particularly short-read junctions, BAM files were processed using EASTR (v1.1.0)^43^ with default parameters before transcriptome assembly.

### Transcriptome Assembly

Transcriptome assembly was performed using several tools and configurations to evaluate performance across different data types (short-read, long-read, hybrid) and conditions. Where applicable (StringTie2, StringTie3, IsoQuant) assemblies were executed in both annotation-free (de novo) mode and annotation-guided mode (using the filtered CHESS 3.0.1 reference annotation). The specific assemblers and key configurations tested included:

#### StringTie3 (v3.0.0)

- Short-read (Default): `stringtie -o output.gtf aligned.bam`
- Short-read (Nascent Mode): `stringtie --nasc -o output.gtf aligned.bam`
- Long-read (Default): `stringtie -L -o output.gtf aligned.bam`
- Long-read (Nascent Mode): `stringtie -L --nasc -o output_lr_nasc.gtf aligned.bam`
- Hybrid (Default): `stringtie -o output.gtf --mix short_read.bam long_read.bam`
- Hybrid (Nascent Mode): `stringtie –nasc -o output.gtf --mix short_read.bam long_read.bam`

Flags: `-G reference_annotation.gtf` was added for annotation-guided assembly.

`--rf` was added for stranded short-read libraries.

#### StringTie2 (v2.2.3)

- Short-read: `stringtie -o output.gtf` aligned.bam
- Long-read: `stringtie -L -o output.gtf` aligned.bam
- Hybrid: `stringtie -o output.gtf --mix short_read.bam long_read.bam`

Flags: `-G reference_annotation.gtf` was added for annotation-guided assembly. `- -rf` was added for stranded short-read libraries.

#### Scallop2 (v1.1.2)

- Short-read: `scallop2 -i aligned.bam -o output.gtf`

Flags: `--library_type first` was used for stranded libraries (equivalent to `--rf` in StringTie).

#### IsoQuant (v3.6)

- Long-read (Annotation-free): `isoquant.py --reference reference_genome.fasta --bam aligned.bam --data_type nanopore -o output_dir`
- Long-read (Annotation-guided): `isoquant.py --reference reference_genome.fasta --bam aligned.bam --data_type nanopore --complete_genedb reference_annotation.gtf -o output_dir`

### Assembly accuracy metrics

Assembly accuracy was evaluated against the filtered CHESS 3.0.1 reference annotation using gffcompare (v0.12.6)^44^. An assembled transcript was considered a correct match (True Positive, TP) if it shared the exact same splice junction chain as a reference transcript and its terminal exon boundaries were within 100 bp of the reference transcript’s boundaries. Sensitivity was calculated as TP / (TP + FN), where FN (False Negatives) represent reference transcripts not found in the assembly. Precision was calculated as TP / (TP + FP), where FP (False Positives) represent assembled transcripts not matching any reference transcript. It should be noted that some FPs may correspond to genuine but unannotated transcripts.

Relative percentage changes in sensitivity (Sr) and precision (Pr) between StringTie3 with the nascent flag (Method 1) and a comparison method (Method 2) were calculated as:

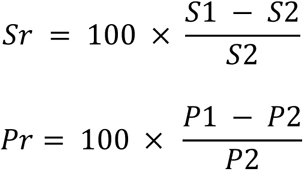

Where S1 and P1 are the sensitivity and precision of StringTie3 (nascent mode), and S2 and P2 are the sensitivity and precision of the comparison method.

### Differential Expression Analysis of Mature and Nascent Transcripts

Transcriptomes were first assembled using StringTie3 (run with `--nasc -G reference.gtf)`. This process generates transcript models in GTF format, where mature and nascent transcripts matching the provided reference annotation are flagged (via `ref_gene_id` and `reference_id` attributes) and include a per-transcript coverage estimate (the `cov` attribute). Gene-level read counts for differential expression analysis were subsequently derived using a custom python script based on this StringTie3 output. Specifically, for each assembled transcript identified by StringTie3 and matching the CHESS 3.0.1 reference annotation, an estimated read count was calculated using the formula:

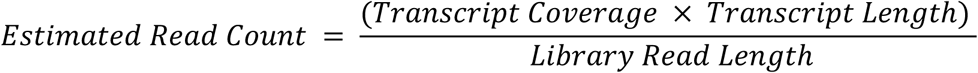

In this formula, ’Transcript Coverage’ corresponds to the value of the `cov` attribute from the StringTie3 GTF output, ’Transcript Length’ is the total exonic length of the assembled transcript in base pairs, and ’Library Read Length’ is the average read length of the sequencing library. For each gene, ’mature counts’ were then calculated by summing these estimated read counts over all assembled transcripts matching annotated mature isoforms of that gene. Correspondingly, ’nascent counts’ were calculated by summing the estimated read counts of assembled nascent transcripts (identified by StringTie3 with the nascent flag and associated with the same parent gene ID) matching that gene.

Differential expression analysis comparing tumor and matched normal samples (breast cancer dataset) or knockout and wild-type conditions (Argonaute dataset) was performed on the derived gene-level counts, separately for mature and nascent transcript populations, using R (v4.3.1) with the edgeR (v4.0.3)^45^ and limma (v3.58.1)^46^ Bioconductor packages. To account for inherent differences between the two RNA fractions, mature and nascent counts were processed independently. Given that low gene expression can lead to incomplete transcript reconstruction by assembly, potentially affecting mature and nascent quantification differently, a stringent filtering step was applied prior to differential analysis. Genes were retained only if they exhibited counts per million (CPM) exceeding 1 in all samples, ensuring analysis was focused on genes with reliable quantification across all conditions; this filter was applied separately to mature and nascent count matrices. Library sizes and compositional biases were normalized using the Trimmed Mean of M-values (TMM) method (`edgeR::calcNormFactors`). Normalized counts were transformed using `limma::voom`, which estimates the mean-variance relationship and computes precision weights for linear modeling (`limma::lmFit`). For the paired breast cancer data, inter-patient variability was modeled using `limma::duplicateCorrelation`, incorporating the resulting correlation into `lmFit` via the block argument (patient ID). For non-paired Argonaute data, standard linear models were used. Contrasts were defined for comparisons of interest, and empirical Bayes moderation (`limma::eBayes`) was applied to stabilize variances and calculate p-values, which were adjusted for multiple testing using the Benjamini-Hochberg method to control the false discovery rate (FDR).

To generate the scatter plots comparing nascent and mature RNA regulation (e.g., Fig. 6), the log₂ fold change (log2FC) values for both the nascent and mature component of each gene were retrieved from the differential expression results described above. For improved clarity and to focus on more robustly expressed genes in the visualization, an additional filter was applied: only genes for which both the nascent and mature components had an average expression (calculated across all samples in the comparison) of at least 8 CPM were included in these plots. In the scatter plots, the log2FC of the nascent fraction was plotted against the log2FC of the mature fraction for each qualifying gene. Genes were considered significantly differentially expressed if the FDR was less than 0.05.

## Data Availability

The RNA-Seq data for DLPFC poly(A) and DLPFC RiboZ libraries are available through the Lieber Institute for Brain Development at http://eqtl.brainseq.org/phase2/ and http://eqtl.brainseq.org/phase1/, respectively. The neuron differentiation long and short read RNA-seq data are accessible in the Gene Expression Omnibus (GEO) under accession number GSE245325, and the breast cancer RNA-seq dataset is available in GEO under accession number GSE103001.

## Code Availability

StringTie3 is implemented in C++ and is freely available as open-source software under the MIT license at https://github.com/gpertea/stringtie. Additional instructions for running nascent mode, including parameters and command-line flags, can be found in the project documentation.

## Acknowledgements

This work was supported in part by NSF grant DBI-2412449 (M.P.) and NIH grants R01-MH123567 and R35-GM156470 (M.P.).

## Author Contributions

I.S. conceived the study, designed and contributed to the implementation of the software, conducted computational analyses, analyzed and interpreted the results, and wrote the manuscript. Z.R. and R.H. aided in the analysis of results. G.P. assisted with software development. M.P. conceived the study, contributed to software, assisted in writing and editing the manuscript, and supervised the entire project. All authors reviewed and approved the final manuscript.

