## Supplemental Information for "StringTie3 Improves Total RNA-seq Assembly by Resolving Nascent and Mature Transcripts"

**Supplementary Table I.** Performance comparison of StringTie2 (ST2) and StringTie3 (ST3, nascent mode) under annotation-guided and annotation-free configurations in rRNA-depleted and poly(A)-selected libraries.

| Library | Assembly Config | Metric | ST2 | ST3 | ∆ | %∆ |
|---|---|---|---|---|---|---|
| rRNA(-) | Annotation-guided | Sensitivity[1] | 38.9% | 38.9% | 0.0% | 0.0% |
| | | Precision[2] | 62.2% | 75.5% | 13.2% | 21.4% |
| | Annotation-free | Sensitivity[1] | 14.0% | 14.2% | 0.3% | 2.1% |
| | | Precision[2] | 32.6% | 36.7% | 4.1% | 12.7% |
| Poly(A)+ | Annotation-guided | Sensitivity[1] | 45.0% | 45.0% | 0.0% | 0.0% |
| | | Precision[2] | 73.0% | 80.6% | 7.6% | 10.5% |
| | Annotation-free | Sensitivity[1] | 17.1% | 16.9% | -0.2% | -0.9% |
| | | Precision[2] | 40.2% | 41.8% | 1.6% | 4.1% |

[1]Sensitivity = TP / (TP + FN), where TP (true positives) are transcripts matching the reference, and FN (false negatives) are reference transcripts missing from the assembly.
[2]Precision = TP / (TP + FP), where FP (false positives) are transcripts not matching the reference.

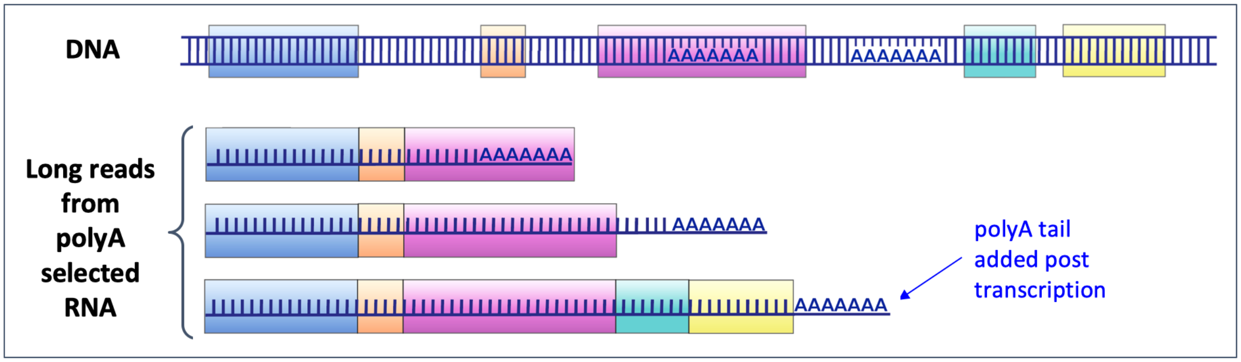


**Supplementary Figure 1.** *Schematic illustrating poly(A) artifacts and nascent intronic poly(A) in long-read, poly(A)-selected RNA-seq libraries.*
The top panel depicts a genomic DNA segment containing exons (colored blocks) and internal poly(A) tracts. Below, three representative long reads from a poly(A)-selected library are shown. The first two reads align to internal poly(A) sequences in the genome, indicating potential artifacts that can arise from priming within genomic poly(A) tracts. The final read illustrates a full-length transcript bearing a genuine post-transcriptionally added poly(A) tail. While poly(A)-selection aims to capture these full-length transcripts, artifacts may arise from internal and intronic poly(A) tracts. StringTie3's long-read module mitigates this issue by identifying and removing reads that map to genomic poly(A) regions, thereby reducing erroneous transcript extensions and improving assembly accuracy.

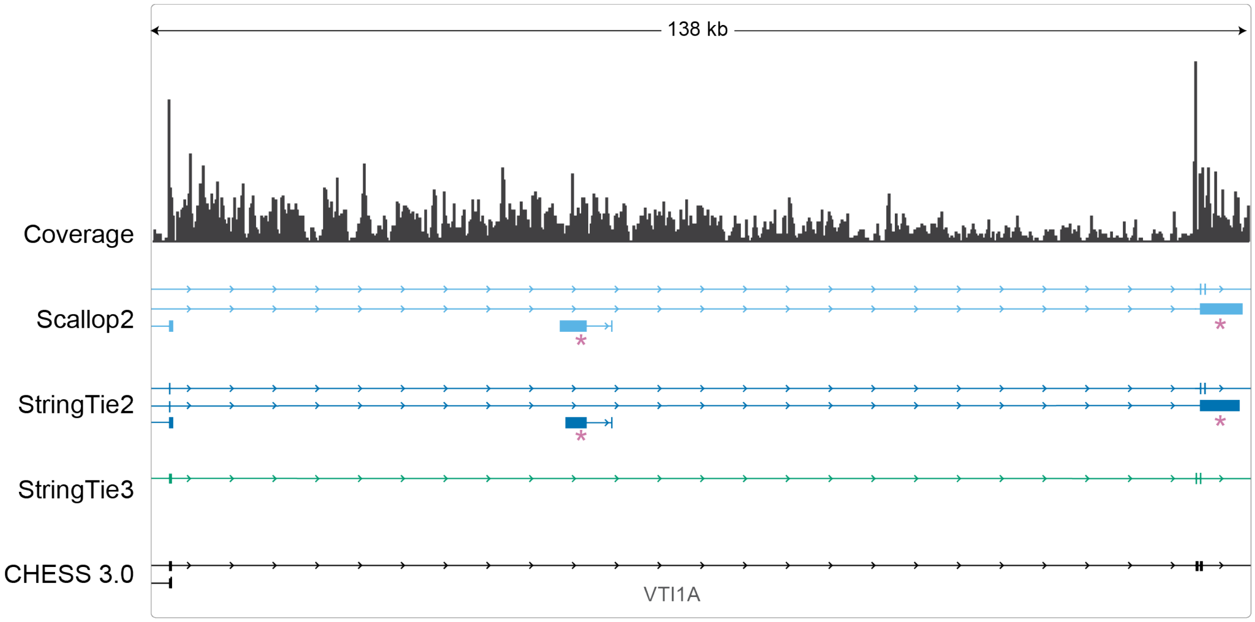


**Supplementary Figure 2.** *Intronic coverage variability results in partial transcript assemblies in Scallop2 and StringTie2, leading to occasional minor sensitivity gain but lower precision*
An example where coverage dips within an intron cause Scallop2 and StringTie2 to produce erroneous transcript assemblies (pink asterisks) that do not match the reference annotation. These dips are misinterpreted as transcript boundaries, resulting in fragmented assemblies. In contrast, StringTie3 classifies these reads as nascent RNA based on the surrounding intronic coverage. One transcript (leftmost, lacking a pink asterisk) is generated by both StringTie2 and Scallop2 and appears in the reference, although low expression and a poly(A) tract at its 3′ end suggest it may be a poly(A)-priming artifact rather than a truly mature isoform. This illustrates how certain reference transcripts sharing intron chains with nascent assemblies can inflate sensitivity for Scallop2 or StringTie2 while reducing precision.

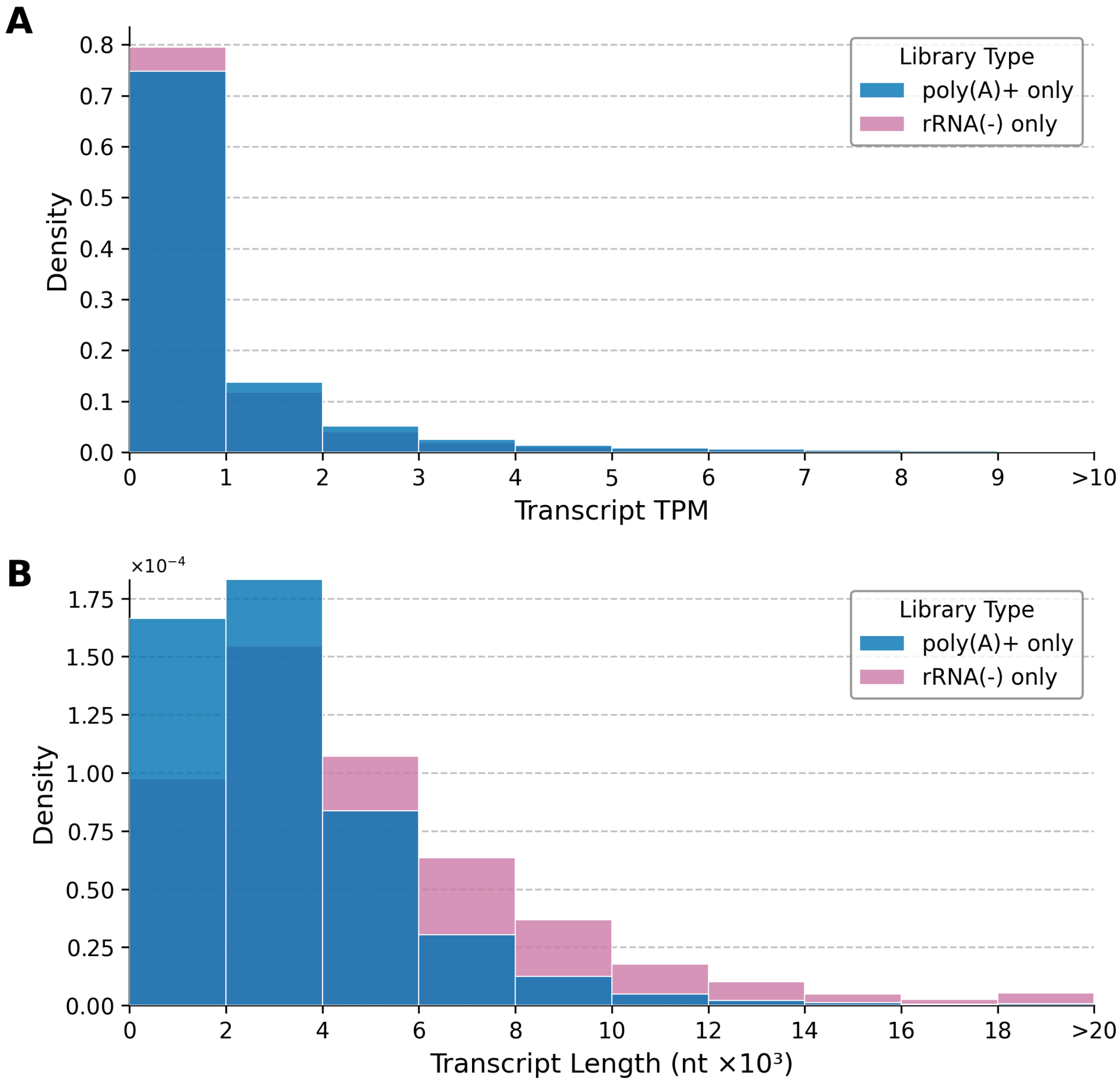


**Supplementary Figure 3.** T*ranscript abundance and length of reference transcripts detected exclusively in rRNA-depleted versus poly(A)+ assemblies across 144 matched DLPFC samples*
**(A)** Distribution of transcript TPM values for reference transcripts detected only in assemblies from rRNA(-) libraries (pink) and only in poly(A)+ assemblies (blue).
**(B)** Distribution of transcript lengths (in nucleotides) for the same mutually exclusive transcript sets (rRNA-depleted-only in pink and poly(A)+ only in blue). The data are aggregated over all 144 paired samples. Transcripts present in both library types are excluded.

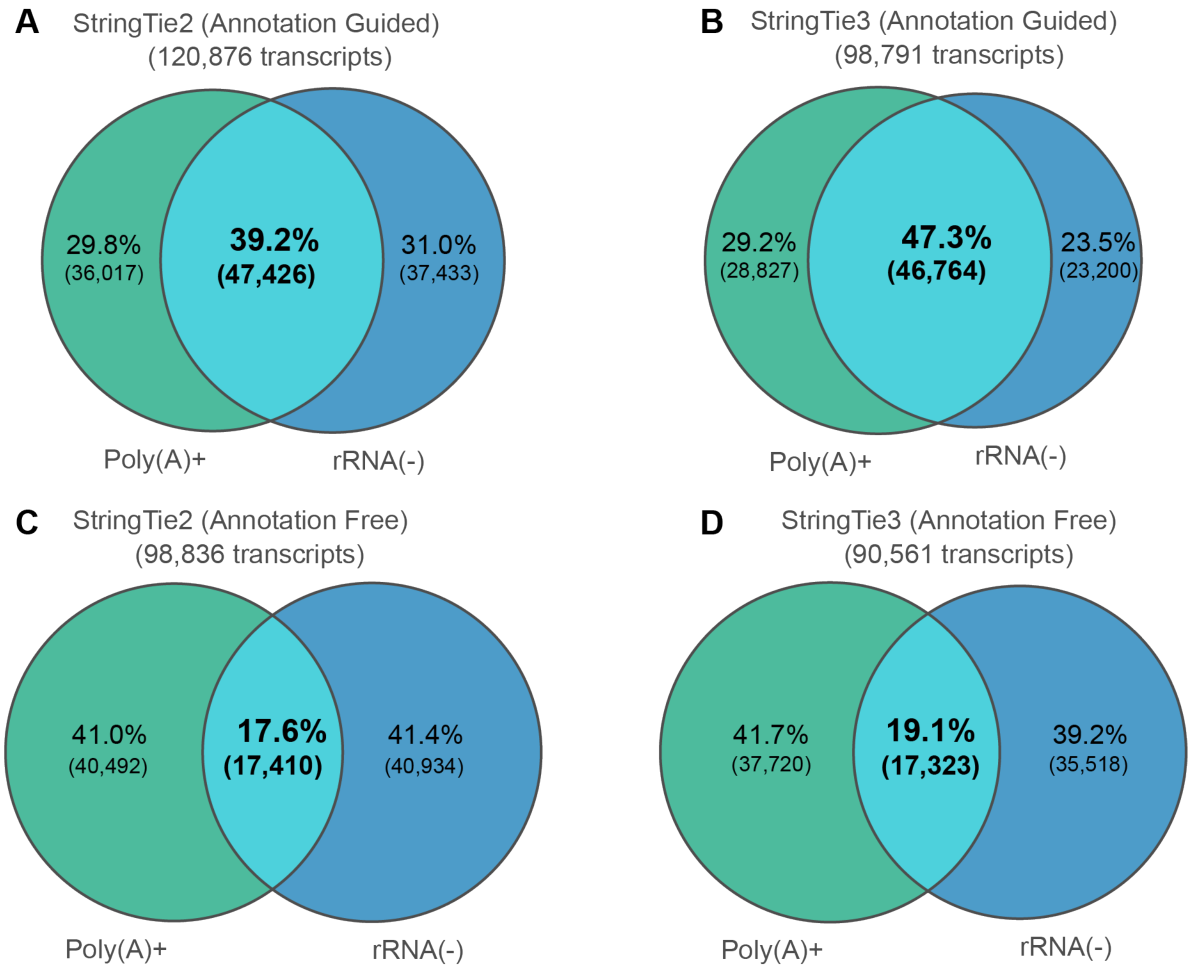


**Supplementary Figure 4.** *Concordance between poly(A)+ and rRNA(–) library assemblies under annotation-guided and annotation-free conditions (StringTie2 vs. StringTie3 with nascent mode).*
Each Venn diagram shows the percentage and count of transcripts classified as either unique to poly(A)+ (green), unique to rRNA(–) (blue), or common to both libraries (overlap). Numbers above each panel indicate the total number of assembled transcripts for that condition (the sum of transcripts from both libraries). All counts and percentages are based on the mean across 144 matched samples**.** Compared to StringTie2, StringTie3 in nascent mode reduces the fraction of transcripts exclusive to one library, while the overlap fraction remains stable or increases, indicating improved concordance across library types.

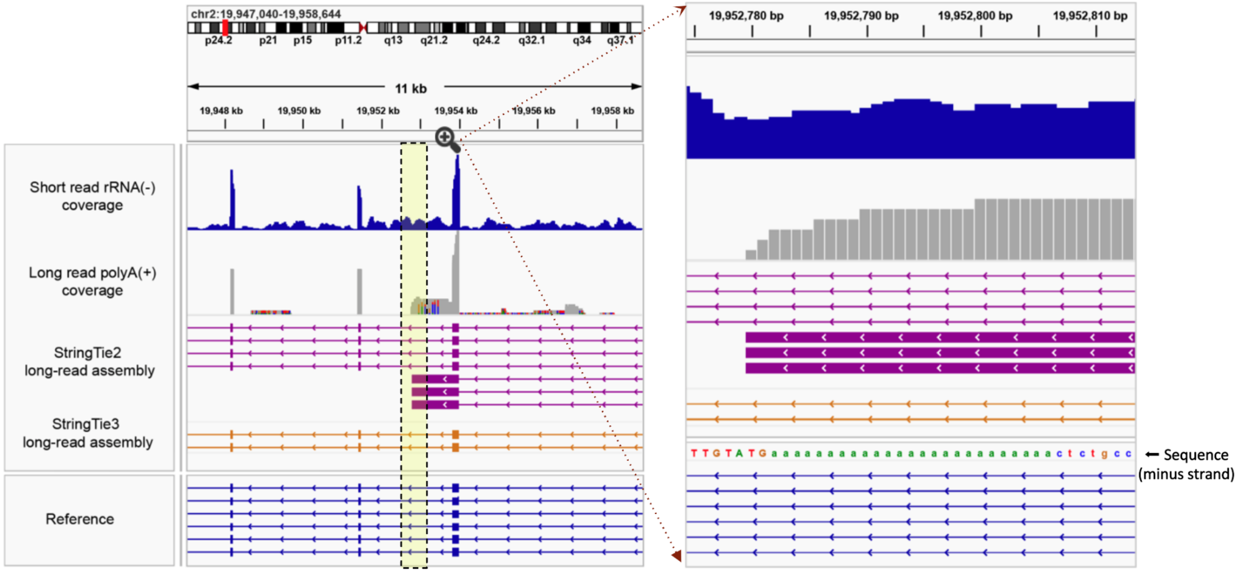


**Supplementary Figure 5.** *Example of intronic priming in a poly(A)+ long-read library and its correction by StringTie3.*

The top panels show short-read rRNA-depleted coverage (blue) and long-read poly(A)+ coverage (gray), with a magnified region indicating long reads that terminate within an intronic portion containing a genomic poly(A) stretch (yellow box). The middle tracks display the resulting assemblies from StringTie2 (purple) and StringTie3 (orange), illustrating how StringTie3 correctly removes intronic poly(A) artifacts that lead to misassembled transcripts in StringTie2. The reference annotation is shown at the bottom (blue).

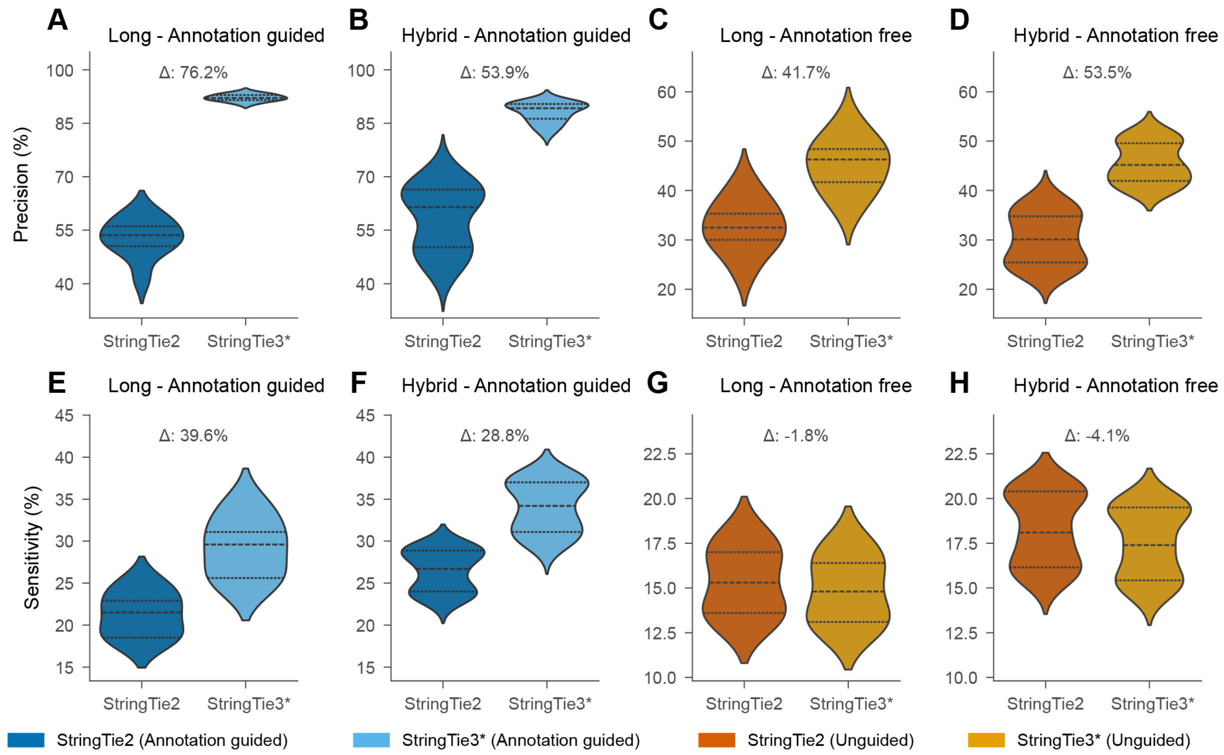


**Supplementary Figure 6.** *StringTie3 demonstrates improved transcript assembly accuracy compared to StringTie2.*

Comparison of transcript assembly performance between StringTie3 with nascent mode (StringTie3* in legend) and StringTie2 across different long and hybrid mode and annotation guided and unguided modes. The figure shows violin plots of precision (A-D) and sensitivity (E-H) metrics from gffcompare analysis. Precision measures the percentage of predicted transcripts that match the reference transcripts, while sensitivity indicates the percentage of reference transcripts that are correctly assembled. The percent change (Δ) of StringTie3 over StringTie2 is shown in each panel. The tighter precision distributions observed in annotation-guided mode in panels (A) and (B), indicate that many samples show precision gains well above the average.
